# Air-liquid interface culture of midbrain organoids improves neuronal functionality and integration of microglia

**DOI:** 10.1101/2023.10.10.561672

**Authors:** Sara Kälvälä, Polina Abushik, Antonios Dougalis, Tarja Malm, Anssi Pelkonen, Šárka Lehtonen

**Affiliations:** A. I. Virtanen Institute for Molecular Sciences, University of Eastern Finland, 70210 Kuopio, Finland; Neuroscience Center, University of Helsinki, 00014 Helsinki, Finland

**Keywords:** Midbrain organoids, air-liquid interface, iPSCs, microglia, Parkinson’s disease, electrophysiology

## Abstract

Midbrain organoids derived from human induced pluripotent stem cells have emerged as a promising in vitro model for studying the molecular and cellular mechanisms in Parkinson’s disease. However, the absence of microglia and the development of a necrotic core in mature organoids have remained key limitations on their utility as functional models of the human midbrain. Here we propose a novel methodology for extended cultivation of midbrain organoid slices, which involves the incorporation of microglia and utilizes an air-liquid culture system. Compared to conventional whole organoids, we found that our model increased the efficacy and consistency of the integration of microglia progenitors. Using single-cell RNA sequencing, we showed that organoid slices maturated in ALI give rise to astrocytes and oligodendrocyte progenitors. Furthermore, the cultivation in air-liquid interface greatly improved neuronal functionality in response to stimulation by NMDA. This stimulation consistently induced robust and stable network activity evaluated by microelectrode array recording. Our newly developed midbrain organoid model with its improved cell type composition and neuronal functionality has potential applications in basic research of Parkinson’s disease and in the development of therapeutics.

**Graphical abstract:** 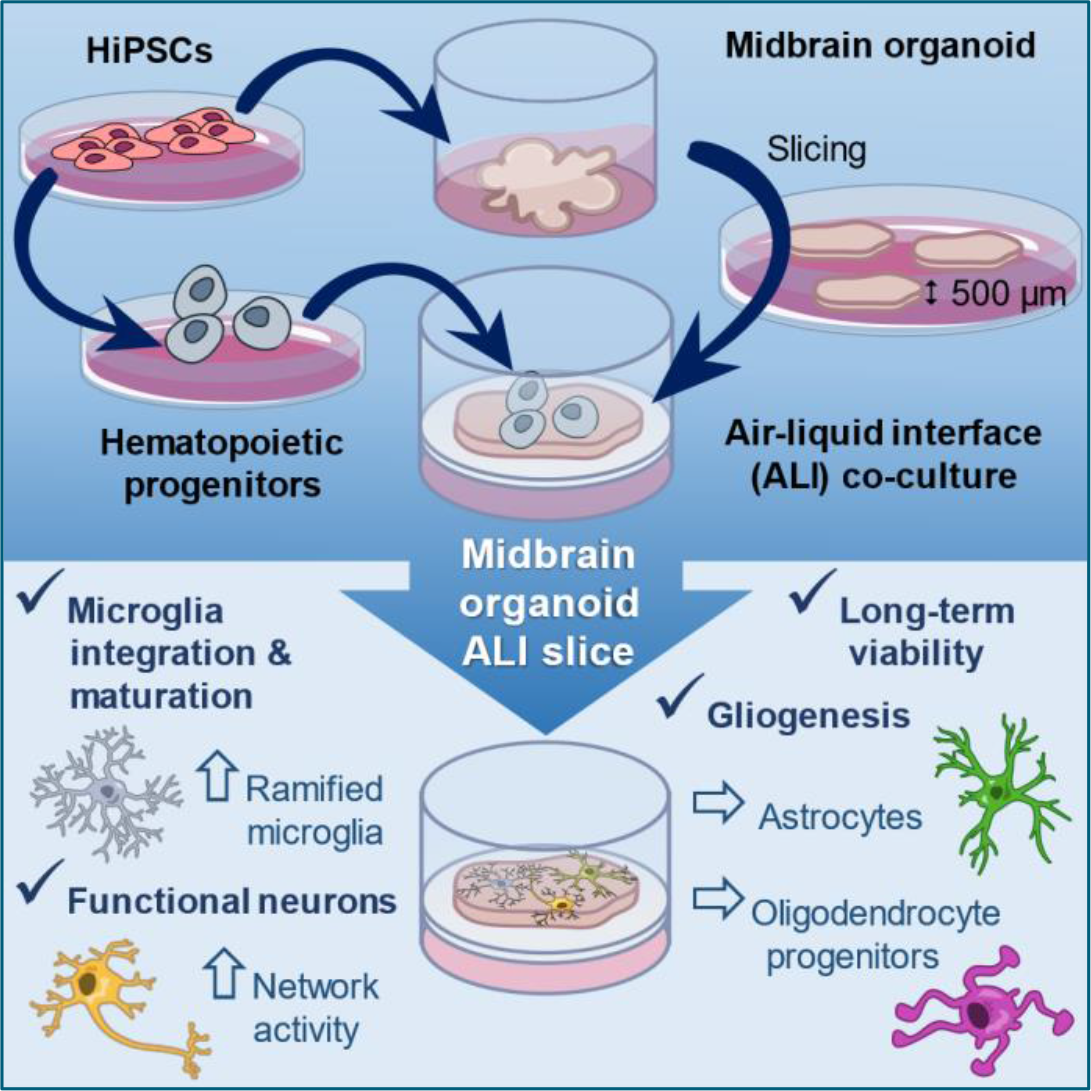

## Introduction

Parkinson’s disease (PD), a neurodegenerative movement disorder caused by the progressive loss of dopaminergic neurons in the midbrain substantia nigra, is increasing in prevalence around the world[1]. PD belongs to a class of neurodegenerative disorders called synucleinopathies, which are defined by the accumulation of aggregated alpha-synuclein protein in the neurons and the glia. It is unclear what initiates the disease process. Most PD cases are sporadic and approximately 5-10% have a known genetic cause[2]. Indeed, some of the most common mutations associated with familial PD have been found to influence the function of glial cells[3]. The most common familial PD locus *LRRK2* has been linked with immune function and microglia[4]–[6]. Subsequently, neuroinflammation and other non-cell autonomous mechanisms have become important areas of research. However, the lack of relevant PD models remains a major limitation. Although animal models have contributed significantly to our understanding of PD biology, the majority of compounds with positive preclinical results do not succeed in clinical trials, which has prompted the call for new human-based disease models[7].

*In vitro* disease modeling has experienced major revolutions within the last few decades. Human-induced pluripotent stem cells (hiPSCs) have made it possible to derive brain cells – neurons, astrocytes, oligodendrocytes, and microglia – that closely resemble human primary cells. Tissue culture has also seen a shift away from traditional 2D cultures towards more *in vivo*-like 3D cultures termed organoids. These advancements have facilitated the development of more complex and physiologically relevant models of brain diseases. Brain organoids grown from hiPSCs can recapitulate aspects of the cell composition and tissue architecture of the human brain[8], [9] and therefore have garnered interest as an alternative to *in vivo* models. Organoids can be grown in large quantities and maintained in culture for long periods of time, making the longitudinal studies of living human brain-like material more accessible and compatible with high-throughput pipelines used in toxicological screening and drug discovery[10], [11]. Still, certain advancements in culture methods are needed to improve the functionality and cell-type composition of organoids to make them more representative of the human brain[12].

Midbrain-like organoids (mORGs) have shown potential as an *in vitro* disease model of PD[11], [13]–[16] but intrinsically lack microglia. Midbrain specification of hiPSCs is achieved by applying exogenous small molecule inhibitors and activators, which drive ectodermal differentiation but inhibit endodermal and mesodermal specification. During development, mesodermal erythromyeloid progenitors migrate into the CNS, where they differentiate into microglial cells[17]. Finding ways to integrate microglia into brain organoids is crucial for faithfully recapitulating the disease processes within an *in vitro* model system, owing to their critical involvement in developmental and neurodegenerative processes. Although several strategies for generating ‘immune-competent’ brain organoids have been described previously[18]– [23], we encountered difficulties in achieving efficient and controlled integration of microglia. As shown in figure 2 panel G, we found that suspension coculture of microglia progenitors and mORGs did not result in sufficient integration outcomes. Instead, we observed that Iba1+ cells were sparsely distributed, mainly residing at the surface of the organoid.

Additionally, we experienced challenges during the implementation of acute microelectrode array (MEA) recording to measure neuronal activity in mORGs. The recordings obtained from whole-organoids yielded minimal activity in response to pharmacological stimulation with N-methyl-D-aspartic acid (NMDA)[12]. Conversely, acute slices of large organoids displayed variability among slices derived from the same organoid. To address these issues, we developed an air-liquid interface (ALI) culture system of mORG slices inspired by the organotypic culture method employed for primary tissue sections[24], [25]. The ALI-cultured slices exhibited consistent and powerful responses to stimulation, resulting in highly synchronous network bursting across nearly all electrodes that were in contact with the sample. Ultimately, the ALI approach enabled more controlled incorporation of microglia-like cells, leading to a homogeneous distribution of Iba1+ cells throughout the tissue. The presence of TH+ neurons was confirmed by immunofluorescent staining. The cell type composition of the slices was studied at 90 and 180 days *in vitro* (DIV) by single-cell sequencing. At 90 DIV neurons were found to be the prevailing cell type. However, by 180 DIV, defined populations of astrocytes and oligodendrocyte progenitors had emerged as the two major cell types. We suggest that the ALI method is superior to the conventional suspension culture of mORGs in terms of facilitating the modeling of non-cell autonomous mechanisms in PD and greatly improving the reliability of functional readouts such as MEA.

## Results

### 1. Air-liquid interface culture of mORG slices facilitates integration and maturation of microglia-like cells

We adapted already established methodology to generate midbrain-like organoids [26]–[29]. Since microglia are highly sensitive to inflammatory stimuli, we used a functionalized silk scaffold as described in[29] to improve nutrient diffusion and prevent necrosis. We confirmed that canonical midbrain marker genes were upregulated during patterning along with the downregulation of the pluripotency gene NANOG (Figure 1, B). Additionally, the co-expression of LMX1A and FOXA2 was confirmed by immunofluorescence (Figure 2, C). To introduce microglia into the mORGs, we derived hematopoietic progenitor cells[30] and co-cultured them in suspension with whole organoids, following a similar methodology as described in reference [18]. The effectiveness of microglia integration into whole organoids was shown to be very low, and the distribution of Iba1+ cells within organoids was discovered to be uneven, with the majority of the cells found near the organoid’s surface (Figure 2, G). In order to address this matter, we implemented a slice culture approach using sections that were 500 μm in thickness and were maintained at ALI (Figure 1, A).

**Figure 1.**
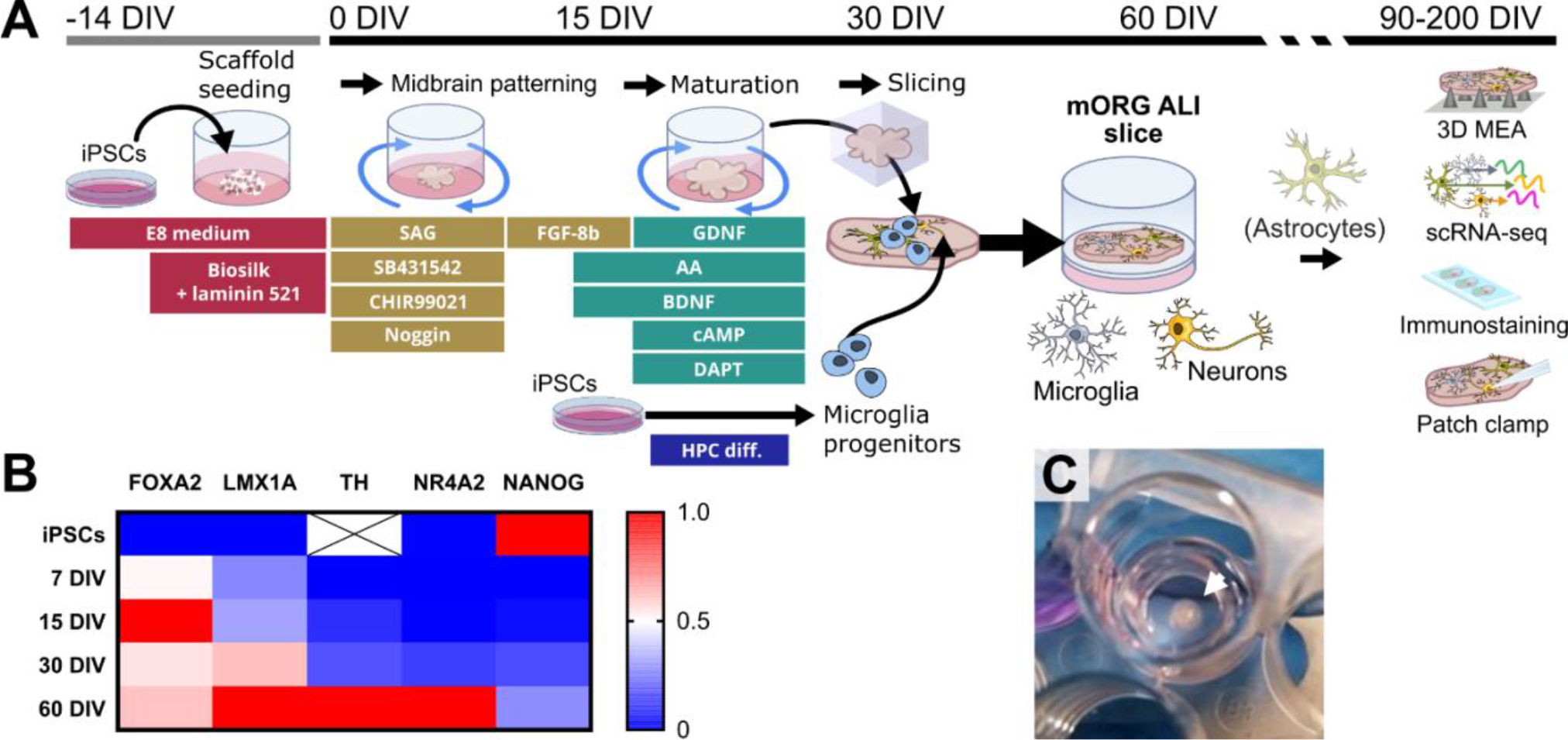
Method overview and validation of midbrain patterning by RT-qPCR. (**A**) Schematic illustration of mORG ALI slice culture workflow with incorporated microglia-like cells. (**B**) Midbrain patterning was validated by RT-qPCR assay in two independent batches. Expression of midbrain markers *FOXA2, LMX1A, TH*, and *NR4A2* increased during differentiation, while the pluripotency gene *NANOG* was suppressed. *TH* was not expressed in iPSCs. Expression values were normalized to *GAPDH*. The color scale is normalized to min-max values. (**C**) Representative image of an mORG ALI slice in culture.

**Figure 2.**
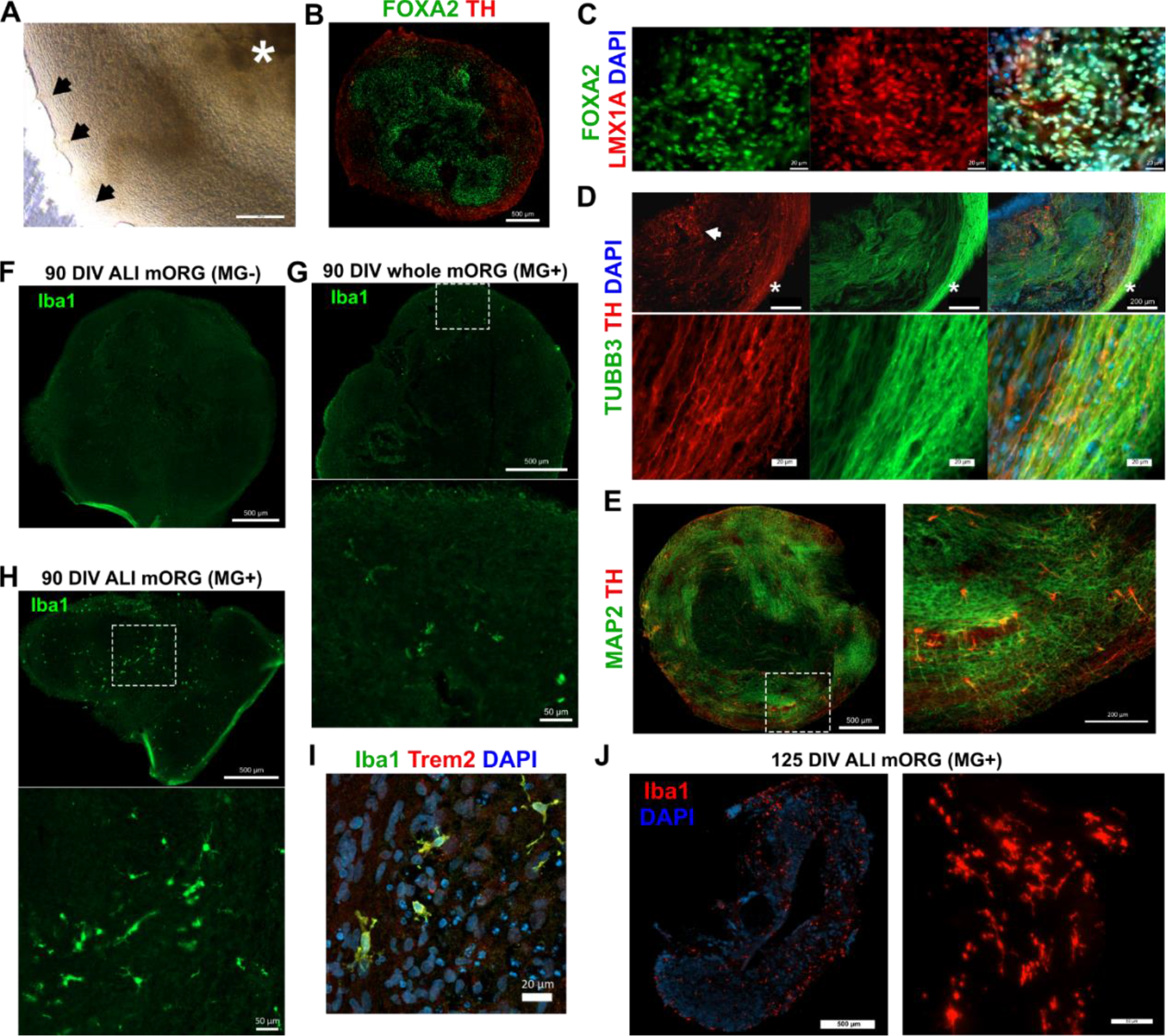
Immunostaining reveals complex cellular morphology in ALI mORGs. (**A**) Brightfield image of a normal mORG slice after 30 days in ALI culture. The edge of the slice appears smooth and clear, with protruding nodules of new growth (black arrows). Silk scaffold fibers are visible in the center of the slice (white asterisk). Scale bar=200 μm. (**B**) FOXA2+ progenitor cells for rosette-like structures in the middle of the slice. Maturing neurons migrate out towards the edge. (**C**) LMX1A and FOXA2 co-localize in the nucleus. Day 40 ALI mORG, scale bar= 20 μm. (**D**) Groups of TH+ neuronal somas are located in the middle layer of the slice (white arrow). A thick bundle of axons wraps around the edge of the slice, as evidenced by intense β3-Tubulin (TUBB3) staining. Bottom panels: a zoomed-in image of fine TH+ processes among the axon bundle (*). Scale bars= 200 μm (upper panels), 20 μm (bottom panels). I MAP2 staining shows the orientation of dendrites in ALI mORGs. Bottom panel: zoomed-in image of the dashed box. (**F**) Negative Iba1 staining in ALI mORG with no added microglia. (**G**) Representative image of a whole mORG with microglia. Only a few Iba1+ cells were observed, with most congregating in one area near the organoid’s surface. (**H**) ALI mORG one month after the addition of microglia progenitors. Iba1+ ramified cells are evenly distributed throughout the tissue. **G-H** Bottom panels: zoomed-in image from dashed box. Scale bars= 500 μm, 50 μm. (**I**) Incorporated microglia are positive for TREM2. (**J**) Ramified microglia persist in the ALI slices 3 months after incorporation. The first panel depicts the entire slice, second panel is a higher magnification image showing microglia processes, scale bars= 500 μm, 50 μm.

Upon introduction of slices into ALI, we observed the emergence of new growth appearing as nodules on the periphery of the slice (Figure 2, A). We noted that the slices grew horizontally in diameter but did not become noticeably thicker. During the process of maturation, a striped pattern developed on the extending border. The FOXA2+ progenitors formed rosette-like structures in the central region of the slice. As the development progressed, the neurons expressing TH protein, indicative of maturation, were observed to emerge closer to the periphery of the slice (Figure 2, B). Immunofluorescent staining for TUBB3 and MAP2 demonstrated the presence of neuronal processes that displayed a unidirectional pattern (Figure 2, D and E). The TUBB3 staining was noticeably more intense along the edge of the slice, suggesting that the new outgrowth consists mainly of axons. Long TH+ processes were also detected inside these tracts. The cell somas and MAP2+ dendrites were located closer the middle of the slice as depicted in Figure 2, panel E. We observed an even distribution of ramified Iba1+/Trem2+ microglia-like cells in the ALI slices 2 months after the addition of microglia progenitors at 90 DIV (Figure 2, H and I). Ramified microglia were present still at 125 DIV, 3 months after incorporation (Figure 2, J). In comparison, whole organoids contained few Iba1+ cells at 90 DIV (Figure 2, G).

### 2. Neurons are the most prominent cell type at 90 DIV

To gain a more comprehensive understanding of the cellular composition of the ALI mORG slices with microglia, we performed single-cell RNA sequencing (scRNA-seq) at 90 DIV. After quality control measures, a total of 8172 cells were accepted for further analysis. A collection of 15 communities were discovered by unsupervised clustering (Figure 3, A). The annotation of cell types was carried out based on the expression pattern of canonical marker genes and validated using the SingleR package and human post-mortem midbrain dataset[31] as reference. We found neurons to be the most abundant cell type accounting for 42% of the total population, distributed across four distinct clusters. Neural progenitor cells (NPCs) were the second most prevalent cell type, comprising 29.4% of the population and distributed across three distinct clusters. A tiny proportion (1.5%) of microglia-like cells was successfully captured. Moreover, other cell types, including neural crest-like cells, unspecific mesenchymal cells, and fibroblasts were also identified. The cycling cells segregate into two clusters, representing NPCs and mesenchymal populations (Figure 3, A & B). The NPCs were characterized by expression of *SPON1* and *HES5*, whereas neurons demonstrated elevated expression levels of *CACNA1B*. Microglia were found to exhibit the expression of *AIF1* and *SPI1* (Figure 3, C). The neuronal population had a transcriptional signature that indicated an enrichment of markers associated with inhibitory (GABAergic) neurons, namely *DLX6-AS1, SLC32A1*, and *GAD1/2*. Neuron cluster 3 was enriched for *RALYL*, a marker for excitatory neurons. Despite confirmed expression of TH by immunofluorescence, we could not detect significant numbers of *TH* transcripts in our data. Through a detailed examination of the neuronal clusters, we were able to single out serotonergic neurons expressing *TPH2* and *SLC6A4*. In contrast, dopaminergic neurons (DaNs) were identified by the presence of *DDC, FOXA2*/*LMX1B*, and *VMAT1* (Figure 3, F). The percentage of DaNs accounted for 2% of the total cell population and 5% of the total neuron population.

**Figure 3.**
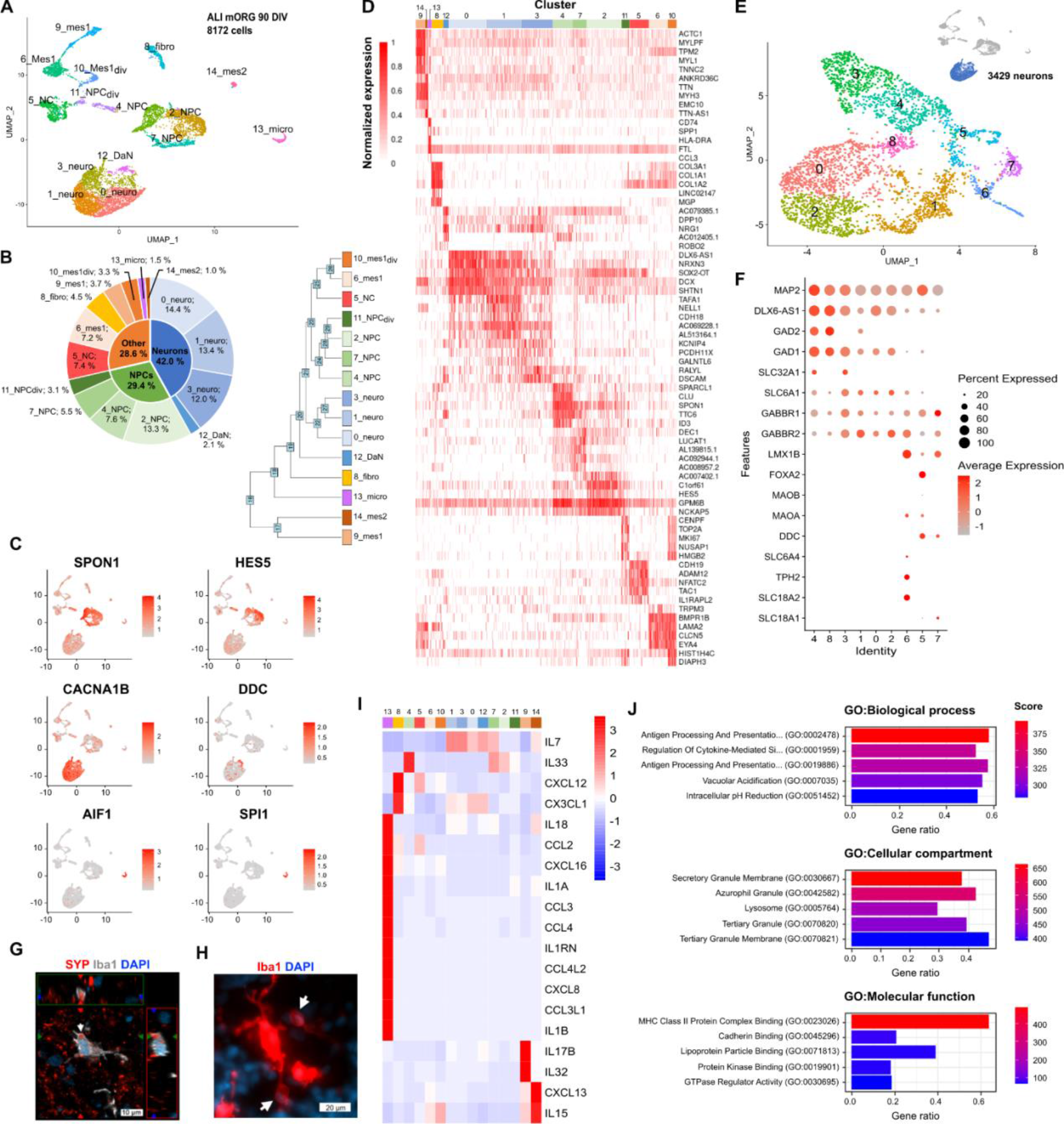
Single-cell transcriptomics of the mORG ALI slice at 90 DIV. (**A**) UMAP embedding of 8172 single cells from 3-month-old ALI mORG. (**B**) Pie chart depicting cell type proportions by category (inner circle), cluster identity (outer ring), and dendrogram depicting cluster similarities. (**C**) Feature plots identifying NPCs, neurons, DaNs, and microglia. (**D**) Heatmap showing expression across all cells for the top 5 differentially expressed genes identified for each cluster and sorted by fold change. (**E**) Neuronal cells were extracted from the data and re-clustered. (**F**) The majority of neurons express genes associated with inhibitory (GABAergic) neurons. Cluster 6 was identified as serotonergic neurons. Dopaminergic marker genes *DDC, LMX1B*, and *FOXA2* are enriched in clusters 5 and 7. (**G**) Confocal image of microglia interacting with synapses (Synaptophysin, SYP). (**H**) Microglia form phagocytic cups engulfing neighboring cell soma. (**I**) Normalized cytokine and chemokine gene expression in each cluster. (**J**) GO-term enrichment analysis of DEGs identified in the microglia cluster. Antigen presenting and phagocytosis-related pathways were upregulated.

We found that the incorporated microglia were interacting with and engulfing synapses and forming phagocytic cups around other cells (Figure 3, G & H). Next, we aimed to investigate if this observed functionality was manifested in the transcriptomic profile. To this end, we performed GO term enrichment analysis using the EnrichR package. We discovered enrichment of antigen processing and presentation, vacuolar acidification and lysosome pathways, implying phagocytic activity (3, J). We also noticed an increase in the activation of cytokine signaling pathways, which prompted us to conduct more in-depth analysis of the mRNA expression levels of cytokines and chemokines across the cell clusters. As expected, our findings indicate that microglia possess the highest expression levels for most cytokines, such as *IL1B, IL1A*, and *CCL2-4*. Neurons were found to mainly express *IL7*, while NPCs had the highest expression of *IL33* (Figure 3, I).

### 3. Astrocytes and oligodendrocyte progenitors emerged by 6 months in culture

We performed scRNA-seq again at 180 DIV to examine the changes in cell composition that occur in aged ALI mORGs. Furthermore, the impact of microglia on the cellular state was investigated. Following the implementation of quality control measures, a total of 3054 cells (mORG-) and 2573 (mORG+) cells were included in the subsequent analysis. The application of unsupervised clustering resulted in identification of 11 distinct clusters of cells. As expected, our findings indicate that glial cells have emerged as the prevalent cell type, constituting more than 60% of cells in both samples. We could identify clusters representing ependymal cells, oligodendrocyte progenitors (OPCs), emerging oligodendrocytes, and astrocytes. Neurons were the minority, making up only 14% of all cells (Figure 4, A-D). Interestingly, distinct grouping of neuroblast-like (NB) cells was exclusively observed in the mORG+ sample. However, we could not detect any microglia-like cells.

**Figure 4.**
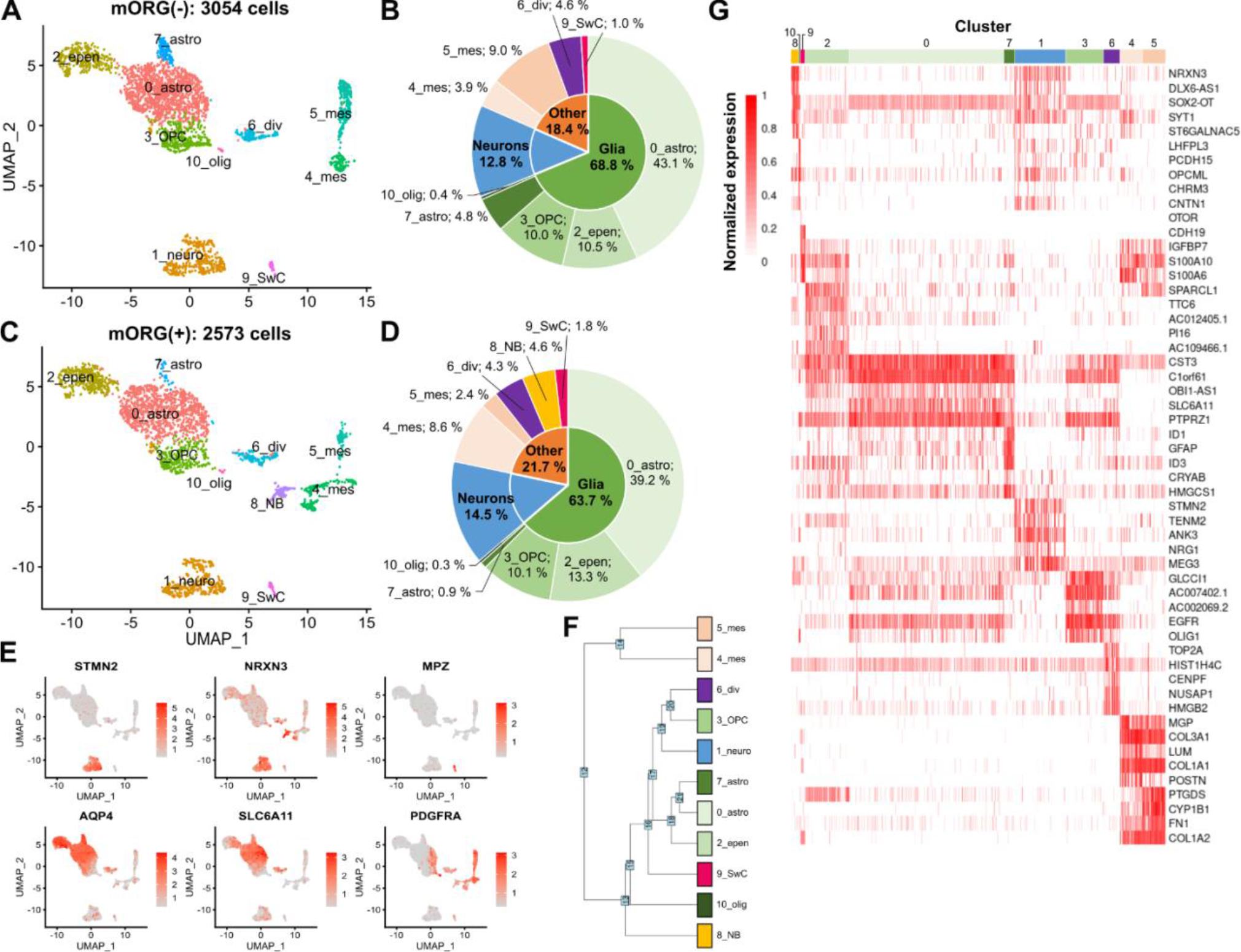
Single-cell RNA sequencing of the mORG ALI slice at 180 DIV. (**A & C**) UMAP projections of single cells from ALI mORG with and without microglia. 11 communities were identified by unsupervised clustering. (**B & D**) Astrocyte-like cells were found to make up most of the cell population. Ependymal cells, OPCs, and oligodendrocyte-like cells were also detected. ALI slices which were enriched with microglia contained a distinct population of neuroblast-like cells, however microglia-like cells were not detected. (**E**) Expression of neuronal genes STMN2 and NRXN3 highlight the main neuronal population and neuroblast-like cells, respectively. Schwann cell-like cells were identified by the expression of MPZ. Both astrocytes and ependymal cells expressed AQP4 at high levels, while astrocytes were the main source of SLC6A11 (GAT3) expression. PDGFRA is expressed by OPCs and early oligodendrocytes. (**F**) Cluster tree analysis. (**G**) Heatmap depicting expression of the top 5 differentially expressed genes in each cluster. Scaled to min-max (0-1).

We noticed that the astrocytes segregated into two clusters, with cluster 7 showing significantly higher expression of *GFAP* and *CRYAB* (Supplementary figure 3, A-B). The proportion of astrocytes belonging to cluster 7 was found to be higher in the microglia null sample (Supplementary figure 3, C). This observation prompted us to further examine the genes that were expressed differentially within this particular population. We discover increased expression of reactive astrocyte markers, and decreased expression of mRNA coding for excitatory amino acid transporters GLT-1 and GLAST-1 (*SLC1A2/3*) which are recognized as markers of homeostatic astrocytes. Furthermore, the expression of *NTN1*, which is involved in axon guidance and has been shown to support survival of substantia nigra DaNs in PD models[32], was also decreased (Supplementary figure 3, D). GO term enrichment analysis indicated upregulation of sterol biosynthesis and glycolytic processes. Genes related to neuronal function were downregulated (Supplementary figure 3, E).

### 4. Air-liquid interface culture increases synchronized network activity

Acute MEA recordings were conducted after 90 DIV using 3D conical electrodes. The organoids were examined in three different formats: whole organoids, acute slices, and ALI cultured slices. Upon stimulation with 200 μM NMDA, we could only detect minimal, if any, induced activity within the intact organoids. The acute slices exhibited heightened response to the NMDA stimulus. We measured two slices from each organoid and noted that the response to NMDA was very heterogenous between the samples. In contrast, in the ALI slices, NMDA consistently induced strong, slice-wide synchronous network bursting (Figure 5, C-E). Additionally, we wanted to find out if exogenous dopamine may potentially affect network activity. We observed that the addition of high concentration of dopamine to the bath resulted in a decrease in the NMDA-induced inter-burst-interval, suggesting the existence of functioning inhibitory dopamine receptors (Figure 5, G).

**Figure 5.**
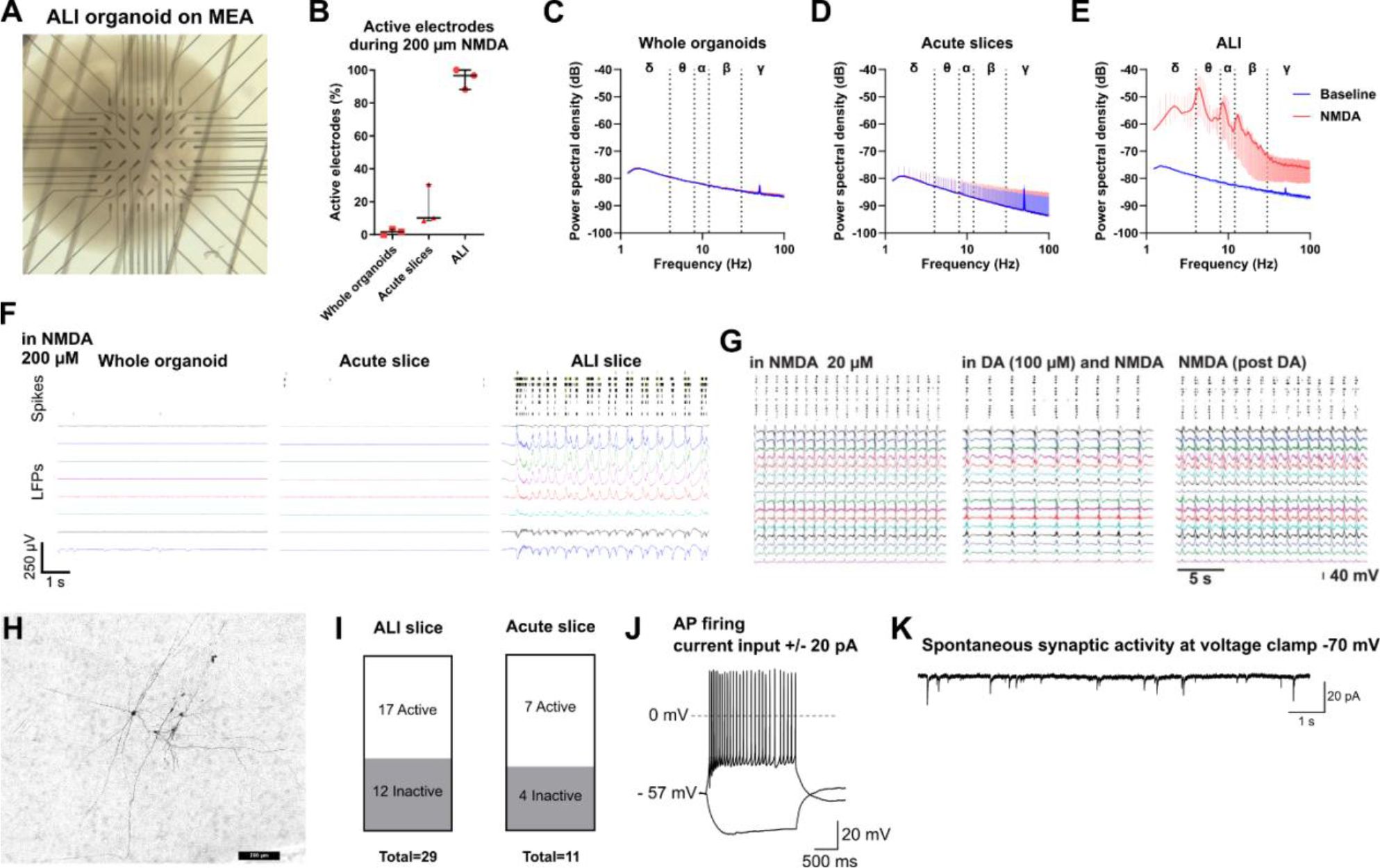
ALI mORGs develop mature and functional networks. (**A**) Representative brightfield image of ALI mORG in MEA well and electrode placement. (**B**) Percentage of active electrodes in different measurement modalities during NMDA stimulation. (**C-E**) Power spectrum density during NMDA stimulation. Measurements from (**C**) whole organoids, (**D**) acute slices, and (**E**) ALI mORGs. (**F**) Representative LFP traces and spike raster from whole organoids, acute slices, and ALI during NMDA stimulation. (**G**) Dopamine suppresses the frequency of NMDA-induced network bursts in ALI mORGs. (**H**) Patched neurons labeled with biocytin and stained by DAB (3, 3’-diaminobenzidine). (**I**) No difference between amount of active and inactive neurons. (**J**) Representative traces showing evoked repetitive AP firing in ALI mORG. (**K**) Representative trace showing spontaneous postsynaptic currents in ALI mORG.

To evaluate functionality of individual neurons in aged ALI mORGs, we performed whole-cell patch-clamp recordings at 186-203 DIV. This time frame was chosen based on our prior research, which suggests that neurons within this age range are likely to display typical electrophysiology activity[19]. The experimental procedure included recordings under voltage-clamp conditions to measure the maximum sodium and potassium currents at specific holding and test voltages (holding voltage – 70 mV, test voltages – 20 mV, and – 10 mV, respectively). Furthermore, the passive leak current density was measured at a holding voltage – 50 mV and test voltage – 120 mV. These measurements served as indicators of neuronal maturation and stage of development as described previously[19], [33], [34]. Recorded currents were normalized to cell size and expressed as picoampere (pA) current per picofarad (pF) of cell capacitance. All values with standard errors for all measured parameters are presented in Supplementary table 1. The cell capacitance and rheobase values were slightly higher in ALI slices compared to acute slices (14.25 [SD 6.71] vs. 9.31 [SD 2.00], p= 0.046 and 26.64 [SD 25.03] vs. 10.86 [SD 5.64], p= 0.036, respectively). The sodium current density was greater in acute slices (-99.18 [SD 102.82] vs. -232.58 [SD 96.12], p=0.009), while hyperpolarization-activated cation (HCN) current density was slightly larger in ALI slices (-6.87 [SD 5.56] vs. -1.03 [SD 2.22]; p=0.008). There was no significant difference observed in the maximal potassium and leak current density.

Incremental voltage steps were applied in voltage-clamp to evaluate the input resistance of the neuron. This was accomplished by subjecting the neuron to a single, 500 ms, -10 mV step from a holding potential of -70 mV. The sodium current density had a direct impact on the frequency of recurrent action potential (AP) firing (Figure 5, J). To investigate the link between current input and spike output firing, as well as the impact of subthreshold currents, the neurons were subjected to incremental current injections of ± 5 pA for a duration of 1 second, starting from their resting membrane potential. The study involved the measurement of various AP characteristics, including threshold, amplitude, and afterhyperpolarization (AHP) amplitude. The AP threshold was defined as the point at which the rate of change of membrane potential (dV/dt) exceeded 10 mV/ms. The rheobase, which refers to the minimum amount of injected current required to elicit spiking from the resting membrane potential, was also determined. As the level of current injection was enhanced, there was a corresponding rise in the cumulative number of APs (Supplementary figure 4, E). The acute slices exhibited a noteworthy rise in the frequency of instantaneous APs when subjected to current injection ranging from 20-45 pA (Mann-Whitney t-test, p<0.05; Supplementary figure 4, F). However, this impact was not maintained as the current was further increased. There was no statistically significant impact observed on the quantity of APs per sweep. Spontaneous excitatory postsynaptic currents (EPSC) were seen in both acute and ALI slices. EPSCs were sampled at a holding potential of -70 mV in 2-minute epochs as reported previously[33], [34].

## Discussion

Previous studies of cerebral organoids [40], [41] and spinal cord organoids[42] in ALI have demonstrated several benefits of the ALI culture, including improved cell survival, reduced hypoxia, and enhanced neuronal functionality. However, this approach has not been implemented before using midbrain organoid slices and incorporated microglia-like cells. Here we show the feasibility of co-culturing microglia progenitors and midbrain organoid slices using the ALI method. Furthermore, we demonstrate that the microglia progenitors readily differentiate into Iba1+ microglia-like cells with ramified morphology. As there are no additional growth factors or cytokines added to the culture medium, we presume that the endogenous factors produced by the organoid are sufficient for inducing microglia specification.

The inclusion of microglia in brain organoid models is crucial for the faithful recapitulation of brain physiology during both normal developmental processes and pathological conditions. Although microglia-like cells can occur spontaneously in unguided brain organoids derived without the use of dual SMAD inhibition, this method introduces uncontrolled variability in the proportion of microglia and the overall cell type composition of the organoids[43]. Guided patterning is preferred for generating more homogenous brain organoids. Still, this approach impedes the spontaneous emergence of microglia in the organoids. Therefore, co-culture approaches using hiPSC-derived microglia or macrophage progenitors have been adopted for introducing microglia to already pre-existing brain organoids. Abud et al. (2017) showed that hiPSC-derived microglia-like cells may effectively integrate into brain organoids. Additionally, they showed a morphological shift from ramified to ameboid in response to a mechanical insult[22]. We also observed a similar morphological change in microglia in ALI slices which were subjected to 3D MEA recording, pointing to diversity in microglial morphology. Sabate-Soler et al. (2022) created assembloids of midbrain organoids and induced macrophage progenitors. They found increased cytokine and chemokine release and increased neuronal excitability in the assembloids[18]. However, the morphology of the microglia appeared mostly ameboid or only partially ramified. This could be due to the fact that the macrophage progenitors were co-cultured with midbrain organoids for only 20 days, while in our study we followed microglia in mORG ALI slices for up to 180 days. It should also be noted that both Abud et al. and Sabate-Soler et al. used substantial numbers of microglia per organoid, specifically 500 000 and 186 000, respectively. The co-culture studies conducted here involved the use of 100 000 microglia progenitors per whole organoid suspended in mORG medium. The outcome of these experiments resulted in uneven distribution of Iba1+ cells with variable numbers of successfully integrated cells. In contrast, 10 000 cells per ALI slice produced an even distribution of Iba1+ cells. Although the supply of hiPSC-derived microglia is not strictly constrained, generating large numbers of these cells requires significant resources. Consequently, this may limit the widespread availability of the technology and its effectiveness in high-throughput experiments. Therefore, the use of ALI slices instead of intact organoids may be preferred due to improved organoid viability, neuron function, and the increased control over microglia incorporation.

A notable shift in cell type composition was found when comparing organoids at 90 DIV and 180 DIV, with glia becoming enriched at the later timepoint. This was expected, as neurons arise early on in development, but astrocytes and oligodendrocytes require longer periods of maturation. The choice of timepoint for experiments should take this shift into consideration. Despite enrichment for CD11b+ cells in the 90-day sample, we only captured a minor population of microglia cells. Enrichment was not conducted for the 180-day sample, resulting in the absence of captured microglia. The proportion of microglia in a single-nuclei RNA-seq dataset obtained from postmortem human midbrain was reported to be approximately 10%[31]. However, the proportion of microglia reported in scRNA-seq studies of embryonal midbrain tissue was around 0.5-1%[44], [45], which aligns more closely with our findings at 90 days. The potential reason for the absence of microglia detection at the 180-days timepoint could be attributed to the limited presence in the midbrain slice and the insufficient number of sequenced cells. We also did not observe major differences in neuronal gene expression between samples with or without microglia, however, microglia may play a role in the regulation of astrocyte homeostasis. In certain cases, astrocytes may adopt a phenotype resembling that of immune cells to compensate for the absence of microglia. We noted enrichment of a reactive-like astrocyte population in ALI slices without microglia.

Despite only minor differences in patch clamp recordings of individual neurons from ALI slices and acute slices, we discovered that the ALI method significantly improved neuronal functionality as evaluated by MEA. At 3 months, intact organoids show almost no response to NMDA. Acute slices of whole organoids improved the response slightly but introduced significant diversity from slice-to-slice. In contrast, the ALI slices, consistently responded with synchronous slice-wide network bursting, which could be modulated by exogenous dopamine. We also discovered that, despite the use of the silk scaffold, whole aged organoids tended to accumulate necrotic tissue in the core. Dead material would slough off during slicing process, making the sections fragile and difficult to handle. Tissue integrity of ALI slices was preserved, making them easier to handle during the recording.

## Conclusions

The novel ALI mORG slicing model presented here offers a promising platform for investigating the cellular and molecular mechanisms in PD. The slicing of the mORG at an early stage allows for homogenous integration of microglia, resulting in enhanced neuronal network activity and functionality. Furthermore, the coexistence of microglia and astrocytes provides an opportunity to explore microgliosis and astrocyte reactivity, as well as their potential interplay in contributing to the progressive degeneration or initiation of dopaminergic neuron loss. The presence of oligodendrocytes, a cell type that only recently gained attention in the PD field, extends the potential for studying inflammatory signaling known to promote neuronal cell death and disease pathology. Moreover, the implementation of a tailored strategy, wherein the cells of the patients themselves are utilized, can facilitate genetic and phenotypic investigations pertaining to PD. Overall, our model provides opportunity to study non-cell autonomous mechanisms and identify novel therapeutic targets for the development of immunomodulatory treatments for PD.

## Materials and methods

### HiPSC culture

HiPSC lines from healthy donors (Cellartis® Human iPS Cell Line 7 (ChiPSC7), Takara Bio, Y00275; MAD8-tdTomato, characterized in[35]) were grown on Matrigel (Corning, 356231) coated dishes and fed daily with Essential 8 medium (Life Technologies, A1517001). The cells were passaged using 0.5 mM EDTA (Life Technologies; cat. 15575) in DPBS (EuroClone, ECB4004L).

### Derivation of midbrain organoids and air-liquid interface slice culture

HiPSCs were dissociated with Accutase (Biowest, L0950) into a single cell suspension. 1.2 million cells in <25μl volume were then added to a vial of Biosilk 521 (Biolamina, BS521-0101) containing 10 μM ROCK inhibitor Y-27632 (Selleckchem, S1049). Silk-cell suspension (7-12 μl) was pipetted to the bottom of a 24-well plate (Sarstedt, 83.3922500) and foamed by pipetting up and down 23 times. The foams were stabilized by placing them in an incubator set at 37 °C for 20 min. The wells were then carefully filled with Essential 8 medium. The hiPSCs were allowed to populate the silk scaffold for 1 week before the scaffolds were detached from the wells and kept on an orbital shaker set at 50 rpm.

To initiate midbrain patterning, the seeded scaffolds were cultured for 7 days with daily medium changes in medium I, containing 10 μM SB431542 (Selleckchem, S1067), 150 ng/ml rhNoggin (Peprotech, 120-10C; SinoBiological, 10267-HNAH), 2 μM SAG (Sigma, SML1314) and 1.5 μM CHIR99021 (Axon, 1386). Then, the medium was switched to mORG medium II, including 100 ng/ml FGF 8b (Peprotech, 100-25). On day 11, 20 ng/mL BDNF (Peprotech, #450-02) and 200 μM L-Ascorbic acid (Sigma-Aldrich, A4403) were included with FGF-8b. Through days 13 to 28 the organoids were maturated in medium II containing 20 ng/mL BDNF (Peprotech, 450-02), 20 ng/mL GDNF (Peprotech, 450-10), 200 μM L-Ascorbic acid, 500 μM bucladesine (Cayman Chemical, 14408) and 1 μM DAPT (Selleckchem, S2215).

After one month in culture, mature organoids were embedded in low-melting point agarose (Lonza, 50101) and sliced to 500 μm thickness with Campden Instruments 7000smz-2 vibratome. The slices were collected in a bath of cold DPBS, then individually placed on 24-well format tissue culture inserts (Millicell, PICM01250), or in groups of 3-4 slices on 6-well inserts (Millicell, PICM03050). Hereafter, the slices were maintained in basal Medium II without additional factors and fed 3 times a week. Slices were allowed to recover for one week before adding microglia progenitors. Microglia progenitors were derived using the hematopoietic stem cell kit (STEMdiff, 05310). Progenitors were collected on days 10-15 from the culture supernatant. 10 000 cells in 1 μl volume were pipetted directly on top of a slice.

### RT-qPCR of midbrain markers

RNA samples were prepared using the RNAeasy kit (Qiagen, 74104) according to manufacturer’s instructions. Extracted RNA was stored at -70 °C. To generate the cDNA libraries, 500 ng of RNA per sample was diluted in ultrapure water, then incubated with 100 pM random hexamer primer (Thermo Scientific, SO142) and 0.5 mM dNTP mix (Thermo Scientific, R0192) for 5 min at 65 °C. A master mix of RT reaction buffer, Maxima RT enzyme (Thermo Scientific, EP0743), and RiboLock RNAse inhibitor (Thermo Scientific, EO0382) was added to the sample and mixed well, then spun down. Final reaction volume was 20 μl. Samples were incubated on a thermal cycler (PTC-100, MJ Research) using the program: 10 min at 25 °C, 30 min at 50 °C, and 5 min at 80 °C. Finally, the cDNA was diluted to 2.5 ng/μl in ultrapure water and stored at -20 °C. TaqMan gene expression assays (Thermo Scientific, Maxima probe/ROX master mix, K0232; *GAPDH*, Hs99999905_m1; *FOXA2*, Hs00232764_m1; *LMX1A*, Hs00892663_m1; *TH*, Hs00165941_m1; *NR4A2*, Hs00428691_m1; *NANOG*, Hs02387400_g1) were run on the StepOnePlus RT-qPCR platform (Applied Biosystems). Expression values were normalized to *GAPDH*.

### Immunofluorescent staining

ALI slices were fixed with 4% PFA for 1 h at 4 °C, then rinsed in 0.1 M phosphate buffer (PB). Tissue was cryoprotected by overnight incubation in 30% sucrose in 0.1 M PB, then embedded in O.C.T. matrix and flash-frozen on dry ice. 20 μm tissue sections were cut with Leica CM1950 cryostat and collected on SuperfrostPlus microscopy slides. Sections were stored at -70 °C prior to staining. All steps of the immunostaining were performed at RT. Sections were washed for 5 min in PBS + 5 min PBST (PBS + 0.05% Tween-20), then permeabilized with 0.4% Triton X-100 for 30 min. The sections were washed again with PBST (3x5 min), then blocked by incubation with 10% normal goat serum (NGS) in PBST for 1 h. Sections were incubated overnight with primary antibodies diluted in 5% NGS (

Table ***2***). Sections were washed with PBST (3x5 min) then incubated with secondary antibodies (Table 3) in 5% NGS for 2 h, followed by 5 min incubation with DAPI (Invitrogen, D1306). Sections were washed 3 x 5 min PBST + 5 min PBS, then mounted with Fluoromount G (Southern Biotech, 0100-01) and sealed with nail polish. Slides were imaged using Zeiss Axio Imager.M2 fluorescence microscope.

**Table 1.** Basal medium components.

| Medium mix | Components | Manufacturer | Catalogue no. |
| --- | --- | --- | --- |
| Medium I | 1:1 DMEM/F-12 | Life Technologies | 21331020 |
|  | 1:1 Neurobasal medium | Life Technologies | 21103049 |
|  | 1:100 MEM-NEAA | Life Technologies | 11140-035 |
|  | 1:200 GlutaMax | Life Technologies | 35050038 |
|  | 1:200 N2 supplement | Life Technologies | 17502001 |
|  | 1:200 Penicillin-streptomycin | Life Technologies | 15140122 |
|  | 50 µM β-mercaptoethanol | Sigma-Aldrich | M3148 |
| Medium II | As above | - | - |
|  | 1:100 B-27 without vit. A | Life Technologies | 12587010 |

**Table 2.**
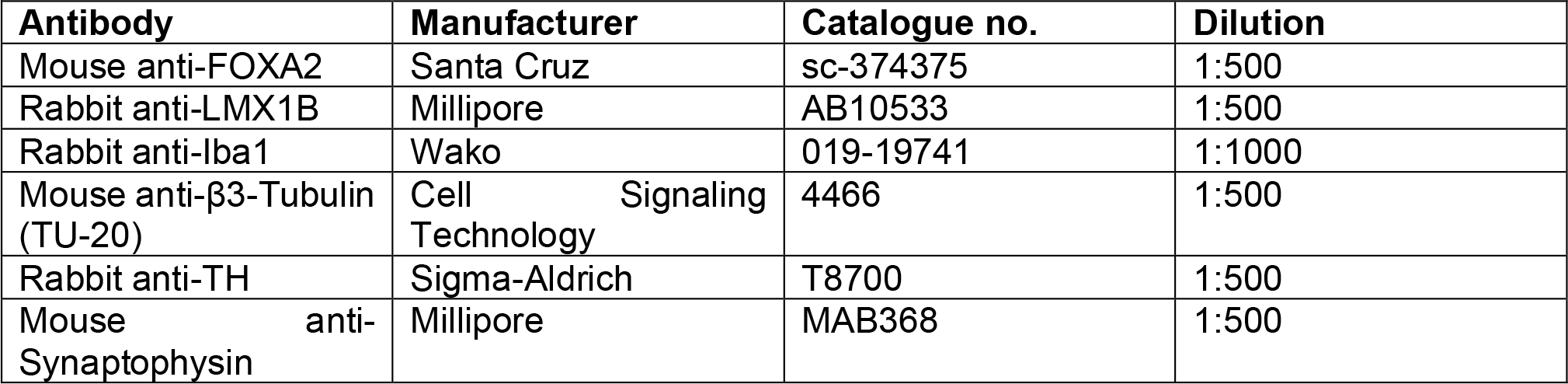
List of primary antibodies used.

**Table 3.** List of secondary antibodies used.

| Antibody | Manufacturer | Catalogue no. | Dilution |
| --- | --- | --- | --- |
| Goat anti-mouse IgG Alexa Fluor 568/488 | Invitrogen | A11004 / A11001 | 1:500 |
| Goat anti-rabbit IgG Alexa Fluor 568/488 | Invitrogen | A11011 / A11008 | 1:500 |

### Single-cell RNA-sequencing

Pooled ALI slices were dissociated into single cells by papain digestion (Worthington, cat.??). Slices were incubated on a shaker at 37 °C for 15 min, then triturated gently using a 1 ml pipette tip. Incubation/trituration was repeated 3 times. The supernatant was collected and centrifuged (300g, 5 min). The pellet was resuspended in ovomucoid inhibitor-containing buffer. Membrane fragments were removed by discontinuous gradient centrifugation (70g, 6 min). Cells were then passed through a 50 μm strainer (Sysmex CellTrics, 04-004-2327). During the preparation of the 90-day sample, we enriched the single cell suspension for microglia by positive selection for CD11b (MACS Miltenyi, 130-049-601; MS columns, MACS Miltenyi, 130-042-201). Half of the cells in the flow-through were added to the microglia fraction. Live/dead staining was done with Trypan blue solution (Sigma-Aldrich, T8154). Dead cells were excluded from the final cell count. Single cell separation was performed on the Chromium Controller platform with Chromium™ Next GEM Chip G (10x Genomics, 1000127). All samples were prepared using the Chromium™ Next GEM Single Cell 3’ Kit v3.1 and Dual Index Kit TT Set A (10x Genomics, 1000269 and 1000215, respectively) according to the manufacturer’s instructions. Library quality was assessed by high sensitivity DNA analysis kit (Agilent, 5067-4626) on the 2100 Bioanalyzer instrument. Library concentration was quantified by Qubit™ dsDNA HS kit (Thermo Scientific, Q32854). Pooled libraries were sequenced on the Illumina NextSeq 500 platform using the High Output Kit v2.5 for 150 Cycles (Illumina, 20024907).

### Sequencing data analysis

Count matrices were generated by processing the raw data on the CellRanger V7 (10x Genomics) pipeline with default parameters and mapping reads to GRCh38-2020-A transcriptome. Downstream analyses were performed using the Seurat package (version 4.3.0)[36] on R version 4.2.2. Low-quality cells with less than 500 genes (empty droplets) or more than 10,000 genes (doublets) or cells with over 10% of reads mapping to mitochondrial genes (dying cells) were filtered out. The counts were log-normalized and top 2000 variable features were identified by vst. Dimensionality reduction (PCA) was applied, and the cells were clustered using Louvain algorithm at resolution 0.3, then visualized by uniform manifold approximation and projection (UMAP). Differentially expressed genes between clusters were identified by Wilcoxon rank sum test (Seurat FindAllMarkers). Genes were considered significantly differentially expressed when adjusted p-value < 0.05. Gene ontology enrichment analysis was performed using the enrichR package[37]. Mitochondrial and ribosomal DEGs were filtered out and pathways with less than 15 annotated genes were excluded. Differentially expressed cytokines and chemokines (HUGO gene groups 601, 598, and 483) were identified and plotted as averaged expression per cluster (dittoHeatmap, dittoSeq R package[38]).

### 60-3D-Microelectrode array recordings

Measurements were obtained and analyzed as previously described[19]. In short, slices were detached from the insert membrane by gently flushing with PBS. Alternatively, the membrane was cut from the insert with the ALI slice attached. The slice was positioned on the electrode array using a brush and a nylon mesh anchor was used to immobilize the slice. Stock solutions of dopamine HCl (Merck, H8502) were made on the day of recording.

### Whole-cell electrophysiology

The brain organoid (186-203 DIV) was transferred and secured with a slice anchor in a large volume bath under an Olympus BX50 WI microscope (Olympus Corporation, Tokyo, Japan) equipped with differential interference contrast (DIC) optics, an x40 water immersion objective and a charge-coupled device (CCD) camera (Retiga R1, Q-imaging, Teledyne Photometrics, Tucson, AZ, USA). Organoids were continuously perfused at a rate of 3-3.5 ml/min at 32-33°C with a recording solution of the following composition (in mM): 120 NaCl, 2.5 KCl, 25 NaHCO3, 1.25 NaH2PO4, 2 CaCl2, 1 MgCl2, 25 Glucose. Organoids from both phenotypes were examined on a single experimental day with a blind protocol to the experimenter. Whole-cell recordings were conducted on Axopatch-200B amplifier (Molecular Devices, San Jose, CA, USA) using 5-8 MOhm glass electrodes filled with an internal solution containing (in mM) 135 potassium-gluconate, 5 NaCl, 10 Hepes, 2 MgCl2, 1 EGTA, 2 MgCl2, 2 Mg-ATP, 0.25 Na-GTP, 0.5 % Biocytin (pH adjusted to 7.3 with osmolarity at 272-285 mOsm/l). Organoids were allowed at least 30 min of settling time in the chamber before any recordings were attempted. Electrophysiology data were low pass filtered at 1 KHz (4 -pole Bessel filter) then captured at 10 KHz via a Digidata 1440A/D board to a personal computer, displayed in Clampex software (version 10.7, Molecular Devices) and stored for further analysis. Single-cell electrophysiological current- and voltage-clamp data were analyzed in Clampfit (Molecular Devices). Synaptic data were detected and analyzed with Mini Analysis software (Synaptosoft Inc., Decatur, GA, USA).

Brain organoids, from which we have successfully recorded neurons, were only selected for inclusion in our database after careful consideration of experimental notes. Accepted neurons exhibited electrophysiological stability during recordings with a stable series resistance (Rs) of less than and stable input resistance (Rin) with less than 15% change after whole-cell break-in and during recording. Biocytin-filled neurons from recorded organoids were subjected to DAB staining as reported previously[39]. Briefly, organoids were fixed in fresh 4% PFA for 48-72 h at 4 °C and then stored indefinitely in PBS with 0.1% sodium azide at 4 °C until processed.

## Author contributions

S.K. developed the ALI slice culture method, prepared and imaged the IF samples, and conducted the single cell sequencing experiments and data analysis. A.P. conducted MEA experiments and data analysis. A.D. and P.A. conducted patch clamp experiments and data analysis. Š.L. and T.M. directed and supervised the work. S.K. wrote the first draft of the manuscript with help of Š.L. and all authors contributed to the final version.

## Data availability

Raw scRNA-seq data and electrophysiology data are available upon request.

## Acknowledgements

We would like to thank the Biocenter Kuopio services, including the Bioinformatics Center, the Cell and Tissue Imaging Unit, the In vitro and ex vivo electrophysiology core facility, the Stem Cell Center, the Single-cell Genomics center, and the Genome Center of Eastern Finland.

