## supplementary materials for "Air-liquid interface culture of midbrain organoids improves neuronal functionality and integration of microglia"

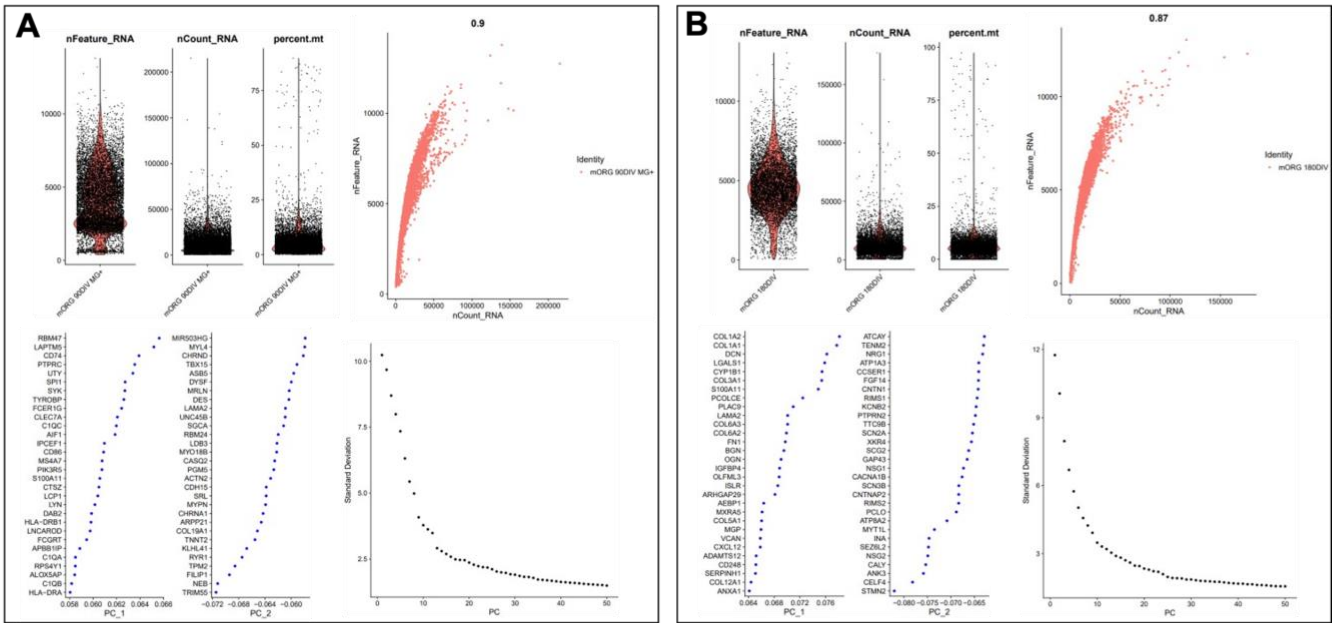


**Supplementary figure 1. Single-cell RNA seq quality control and PCA metrics.** (**A**) ALI mORG 90 DIV dataset. (**B**) ALI mORG 180 DIV dataset.

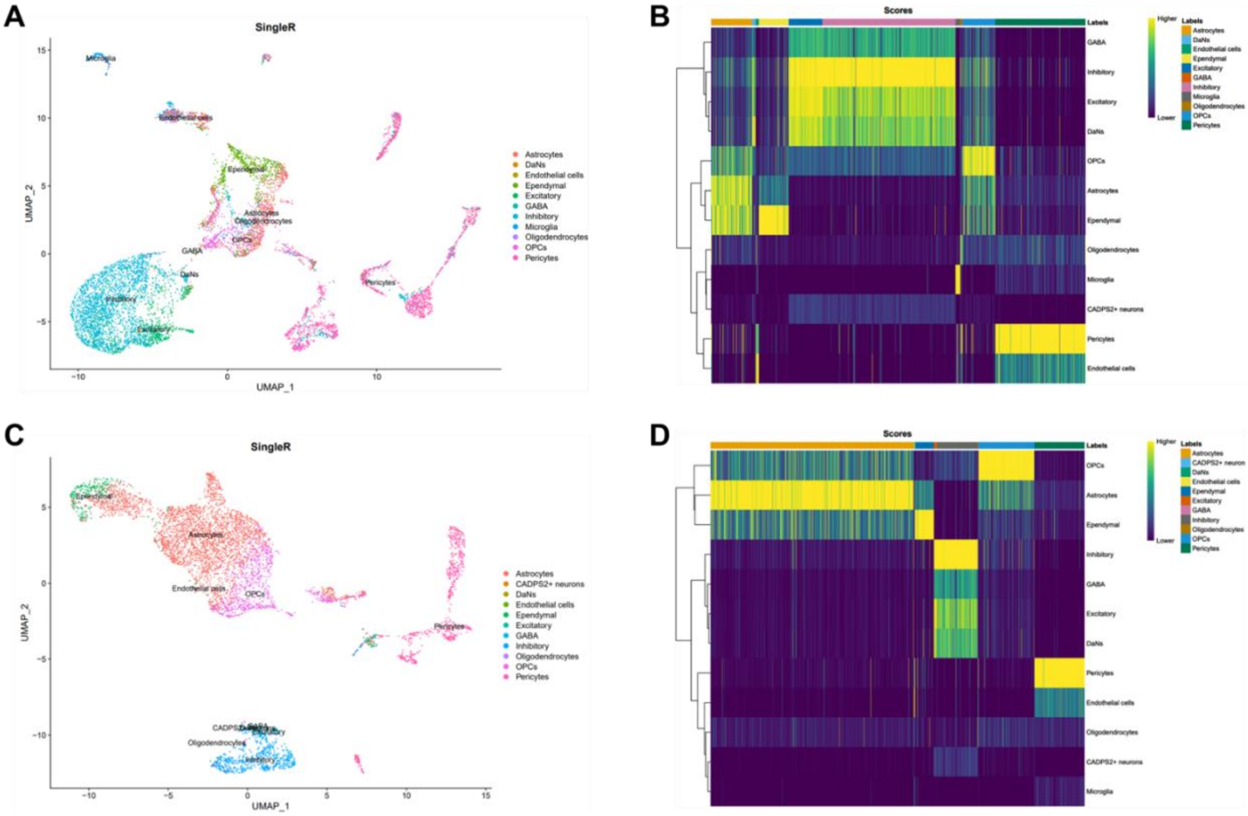


**Supplementary figure 2. SingleR cell ontology annotations and scores.** A dataset of single-nuclei sequencing of human postmortem midbrain tissue[31] (Smajic et al. 2021) was used as the reference. (**A**) Ontology annotations for the 90 DIV mORG dataset and (**B**) heatmap of annotation score per cell. (**C**) Ontology annotations for the 180 DIV mORG dataset and (**D**) heatmap of annotation score per cell.

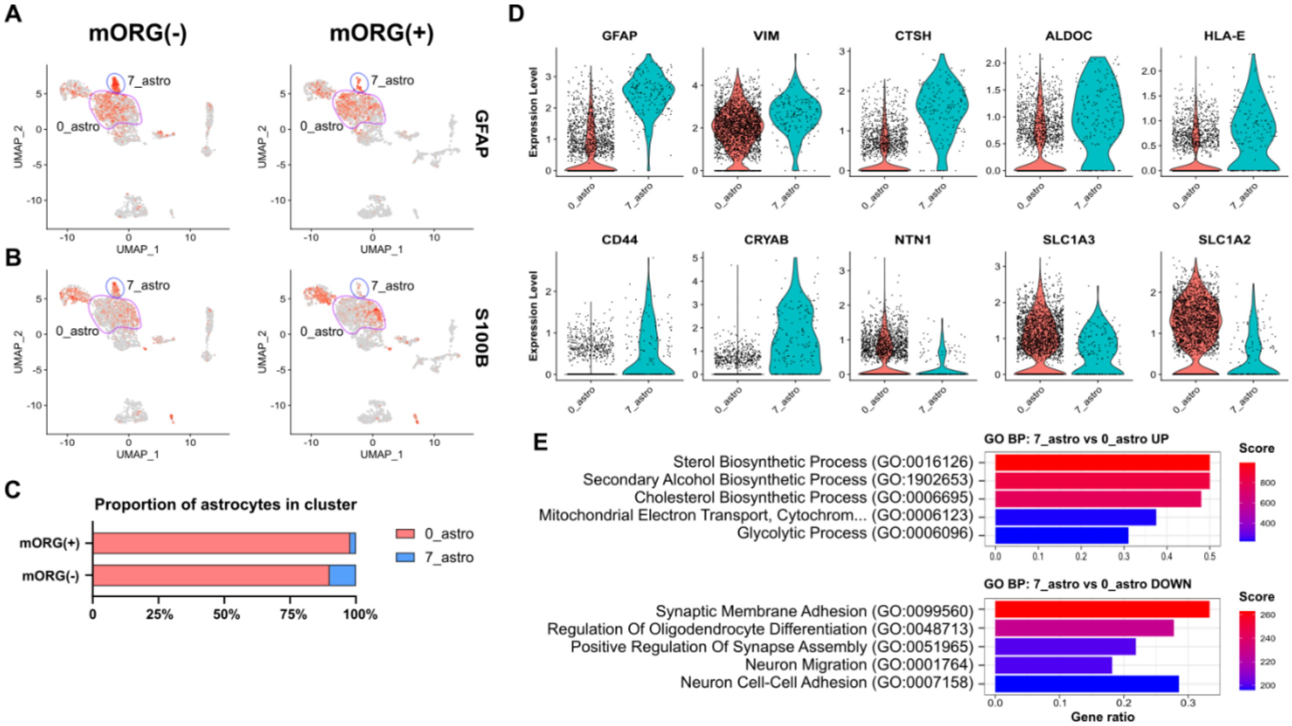


**Supplementary figure 3. Reactive-like astrocyte population is reduced in aged ALI mORGs with microglia.** (**A-B**) Expression of canonical astrocyte markers GFAP and S100B. (**C**) Distribution of astrocytes in each cluster. Reactive astrocytes make up ~10% of all astrocytes in mORG(-) sample, while in mORG(+) sample the frequency is ~2%. (**D**) Violin plots depicting expression of astrocyte activation-associated markers (*GFAP*, *VIM*, *CTSH*, *ALDOC*, *HLA-E*, *CD44*, and *CRYAB*), axon guidance and survival gene *NTN1*, and homeostatic transporter genes *SLC1A3/2* (GLAST-1/GLT1), between clusters 0 and 7. (**E**) GO term enrichment analysis shows upregulation of lipid synthesis and glycolysis in cluster 7 astrocytes, while neurodevelopmental pathways are downregulated. mORG(-), midbrain organoid slide without microglia; mORG(+), midbrain organoid slide with integrated microglia.

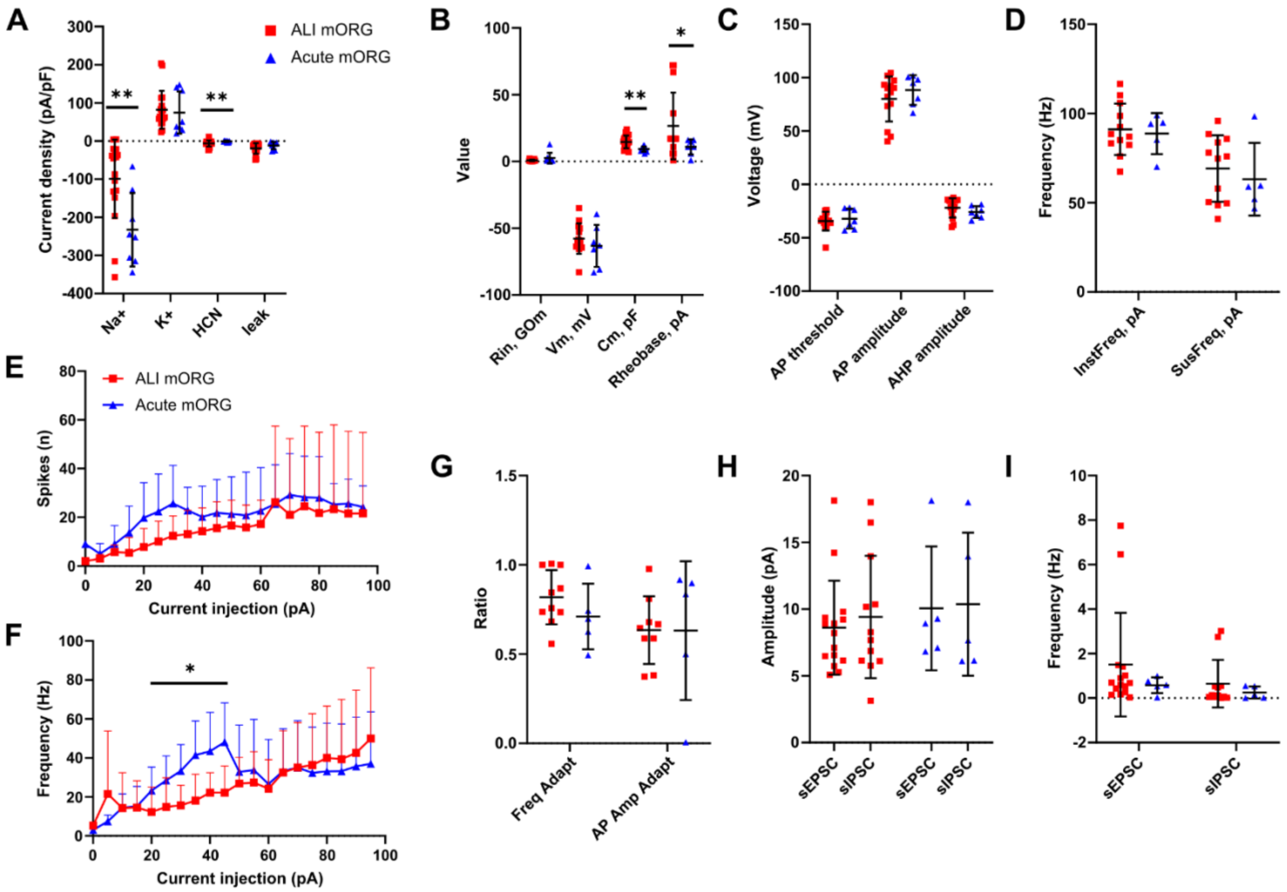


**Supplementary figure 4. Passive and active ionic properties of recorded neurons from acute slices and ALI slices.** (**A**) Current density of sodium, potassium, HCN, and leak channels. * Mann-Whitney t-test, $p<0.05$ (**B**) Input resistance, resting potential, capacitance, and rheobase values. (**C**) AP threshold, amplitude, and AHP amplitude. (**D**) Instantaneous and sustained frequency. (**E**) Spikes per sweep against injected current. (**F**) Instantaneous frequency against injected current. *Mann-Whitney t-test, $p<0.05$. (**G**) Frequency and amplitude adaptation (ratio to baseline?). (**H**) Amplitude of spontaneous excitatory (sEPSC) and inhibitory (sIPSC) postsynaptic currents. (**I**) Frequency of spontaneous postsynaptic currents. All values are given as mean +/- SD. ** Mann-Whitney t-test, $p<0.01$.

**Table 1. Summary of passive and active ionic properties and action potential characteristics of neurons recorded from mORG slices grown in ALI and acute slices of whole mORGs.** Mann-Whitney non-parametric, pairwise comparison with a p-value < 0.05 considered statistically significant.

| | **ALI slice** | | | **Acute slice** | | | **Mann-Whitney t-test** | | | |
|---|---|---|---|---|---|---|---|---|---|---|
| *Parameter, unit* | *Mean* | *SD* | *N* | *Mean* | *SD* | *N* | *p-value* | *Mean rank diff.* | *U* | *sig.* |
| Rin, GOm | 0.84 | 0.59 | 18 | 2.53 | 3.93 | 9 | 0.118 | -5.167 | 50 | |
| Vm, mV | -57.82 | 11.41 | 14 | -63.18 | 15.70 | 7 | 0.400 | 2.571 | 37 | |
| **Cm, pF** | **14.69** | **4.80** | **18** | **9.31** | **2.00** | **9** | **0.003** | **9.250** | **25.5** | ** |
| **Rheobase, pA** | **26.64** | **25.03** | **14** | **10.86** | **5.64** | **7** | **0.036** | **5.893** | **21.5** | * |
| **Na+, pA/pF** | **-99.18** | **102.82** | **18** | **-232.58** | **96.12** | **8** | **0.009** | **8.306** | **26** | ** |
| K+, pA/pF | 81.87 | 49.64 | 18 | 74.48 | 55.34 | 8 | 0.461 | 2.528 | 58 | |
| **HCN, pA, pF** | **-6.33** | **8.18** | **18** | **-1.03** | **2.22** | **8** | **0.006** | **-8.667** | **24** | ** |
| leak, pA/pF | -19.53 | 14.02 | 18 | -11.80 | 9.59 | 8 | 0.144 | -4.875 | 45 | |
| AP threshold, mV | -34.39 | 8.77 | 14 | -32.09 | 9.40 | 7 | 0.913 | -0.429 | 47 | |
| AP amplitude, mV | 80.04 | 21.12 | 14 | 88.33 | 13.95 | 7 | 0.443 | -2.357 | 38 | |
| AHP amplitude, mV | -21.96 | 9.05 | 14 | -25.92 | 5.62 | 7 | 0.197 | 3.857 | 31 | |
| InstFreq (at 50 pA current injection), pA | 91.11 | 14.39 | 12 | 88.71 | 11.52 | 5 | 0.959 | -0.283 | 29 | |
| SusFreq (at 50 pA current injection), pA | 69.17 | 18.63 | 12 | 63.17 | 20.37 | 5 | 0.799 | 0.850 | 27 | |
| Freq Adapt (at 50 pA current injection), ratio | 0.82 | 0.15 | 10 | 0.71 | 0.18 | 5 | 0.197 | 3.300 | 14 | |
| AP Amp Adapt (at 50 pA current injection), ratio | 0.63 | 0.19 | 9 | 0.63 | 0.39 | 5 | 0.699 | -1.089 | 19 | |
| sEPSC Freq, Hz | 1.50 | 2.33 | 15 | 0.58 | 0.35 | 5 | 0.626 | 1.600 | 31.5 | |
| sIPSC Freq, Hz | 0.65 | 1.07 | 12 | 0.25 | 0.27 | 5 | 0.779 | 0.850 | 27 | |
| sEPSC Amp, pA | 8.61 | 3.52 | 15 | 10.05 | 4.64 | 5 | 0.406 | -2.667 | 27.5 | |
| sIPSC Amp, pA | 9.41 | 4.58 | 12 | 10.37 | 5.35 | 5 | 0.859 | -0.567 | 28 | |
